# Extended Quality (eQual): Radial threshold clustering based on n-ary similarity

**DOI:** 10.1101/2024.12.05.627001

**Authors:** Lexin Chen, Micah Smith, Daniel R. Roe, Ramón Alain Miranda-Quintana

## Abstract

We are transforming Radial Threshold Clustering (RTC), an *O*(*N* ^2^) algorithm, into Extended Quality Clustering, an *O(N)* algorithm with several novel features. Daura et al’s RTC algorithm is a partitioning clustering algorithm that groups similar frames together based on their similarity to the seed configuration. Two current issues with RTC is that it scales as *O*(*N* ^2^) making it inefficient at high frame counts, and the clustering results are dependent on the order of the input frames. To address the first issue, we have increased the speed of the seed selection by using *k*-means++ to select the seeds of the available frames. To address the second issue and make the results invariant with respect to frame ordering, whenever there is a tie in the most populated cluster, the densest and most compact cluster is chosen using the extended similarity indices. The new algorithm is able to cluster in linear time and produce more compact and separate clusters.

## Introduction

As the amount of data that can be generated continues to increase largely thanks to advances in hardware, such as the widespread adoption of graphical processing units for molecular dynamics (MD) simulations, it is critical that the methods and algorithms used to analyze this data remain efficient so as to avoid becoming a bottleneck. ^1–3^ The data generated from a MD simulation is typically in the form of a high-dimensional trajectory which can contain a multitude of information such as atomic positions and velocities, forces, etc., and in turn can be used to generate derived data such as coordinate RMSD (root-mean-square deviation) from a reference structure, dihedral angles, secondary structure content, and so on.^4,5^ Although the raw high-dimensional trajectory provides a complete picture of the dynamics of a biomolecule, this is usually impractical for analysis. Hence, it is often useful to employ some sort of dimensionality reduction to only retain the dominating features of a simulation.^6–9^ One such very popular method is clustering, in which conformations or states are classified and grouped together based on some sort of similarity metric. ^10–18^ Cluster analysis can be thought of as unsupervised learning, which itself is a form of machine learning that does not have available labels, but instead classifies the data using only information about its internal structure. Clustering is especially useful in the case of MD simulations, where it can be employed to figure out some of the most representative configurations of a system, as a starting point to build kinetic models, and to aid in virtual screening and free energy perturbation pipelines, just to name a few. ^1,14,18–22^

Non-hierarchical clustering algorithms have been a widely used clustering method for their simplicity, scalability, and efficiency, such as *k*-means,^19,23^ and *k*-medoids. ^24^ Radial Threshold Clustering (RTC) was introduced by Daura and colleagues. ^12,13^ Essentially, it clusters frames in a trajectory that is within the RMSD threshold from the seed (the growing point of a cluster). This is a very popular clustering strategy, with the Taylor-Butina clustering^25^ algorithm sharing the same idea as Daura’s radial threshold clustering (RTC) algorithm but its main applications are found in cheminformatics and it uses similarity metrics specific to cheminformatics, like the Tanimoto index. Another algorithm that has been compared to the RTC algorithm is the quality or diametral threshold clustering. ^26^ Quality Threshold Clustering is a diametric clustering algorithm that groups similar frames based on their similarity to every accepted member of a cluster for every iteration. This algorithm requires two loops; one to iterate every frame to propose its cluster and another one to iterate every member in a cluster to calculate similarity with a candidate which, if not implemented carefully, quickly becomes an *O*(*N* ^5^) time complexity method. ^26,27^Although it preserves the quality of the cluster just like the RTC, the diameter-based approach is certainly far from ideal for bigger datasets.

A common theme between these two techniques is the requirement to pre-compute the RMSD pairwise matrix as the input, which has *O*(*N* ^2^) time and memory complexity. ^28^ Recently, we have proposed a new way to compare molecules and conformations based on combining multiple objects at the same time, that is, using *n*-ary functions, which have a much more attractive *O(N)* scaling. ^29–33^ This technique has been previously applied in sampling, chemical space exploration, dissecting epigenetic libraries, in mass spectrometry studies, and in hierarchical and *k*-means clustering. ^30,33–37^ Here, we propose to leverage the efficiency of the *n*-ary metrics, as well as *k*-means to propose a linear scaling variant of RTC. The key is to use these methods to quickly identify potential cluster centroids, without the need to explore every single conformation. We incorporate our extended quality (eQual) method in our Molecular Dynamics Analysis with *N*-ary Clustering Ensembles (MDANCE) package, which is a software with several clustering modules with applications to not only molecular simulation, but also to cheminformatics, mass mass spectrometry, and other fields.^28,38^ We currently have similarity metrics implemented for cheminformatics and the MD simulations use but additional similarity metrics can be easily integrated into our current framework. In the following sections we introduce MSD as a convenient and efficient metric, discuss the eQual algorithm, and show that it closely matches the RTC results, but without having to generate a memory- and time-intensive pairwise RMSD matrix.

### Theory

#### Radial Threshold Clustering

Introduced by Daura et al, radial threshold clustering is a popular clustering algorithm that partitions the simulation data using a radial threshold to form spherical clusters. ^39^ (The prototypical radial clustering algorithm is represented in Fig. 1.) It starts with a matrix of pairwise RMSD values between all frames. This is already an *O*(*N* ^2^) method. Following the mathematical notation in Daura et al. *x*_*k*_ ∈ *S*_*m*_ represents the seed, or the growing point of a cluster. *S*_*m*_ is your matrix. *S*_*m*_ is a *N* × *N* matrix, where *N* is the number of frames. *S*_*m*_(*i, k*) is the RMSD between frame *i* and *j. A*_*m,k*_(*θ*) = {*x*_*i*_ ∈ *S*_*m*_ : *rmsd*_*ki*_ < *θ*} is the set of frames that are within the threshold distance of the seed, *x*_*k*_. *A*_*m*_(*θ*) = {*A*_*m,k*_(*θ*) : *k* ∈ *I*_*m*_} means for every index, *k* in all indices *I*_*m*_, every available configuration proposes a cluster, *A*_*m,k*_(*θ*). Every cluster, *A*_*m,k*_(*θ*), is compared to other proposed clusters *A*_*m*_(*θ*) and ranked by the size, |*A*_*m,k*_(*θ*)|, in the cluster. The cluster with the most objects is the winner. If there is a tie, the most common settlement is to take the first cluster on the winner list as the ultimate winner. The winner is then removed from the list and the process is repeated until there are no more frames left.

**Figure 1.**
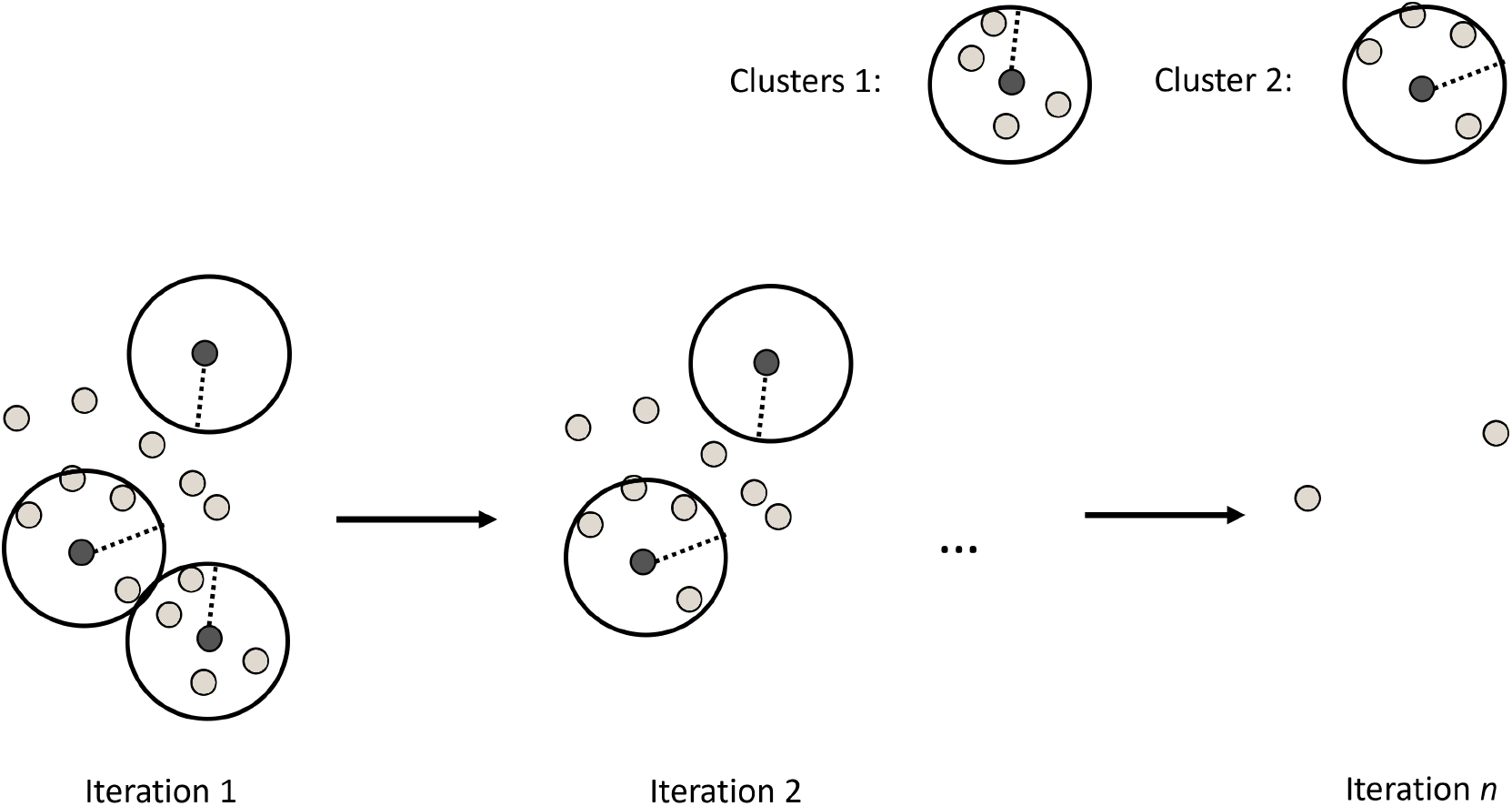
eQual schematics.

**Figure 2.**
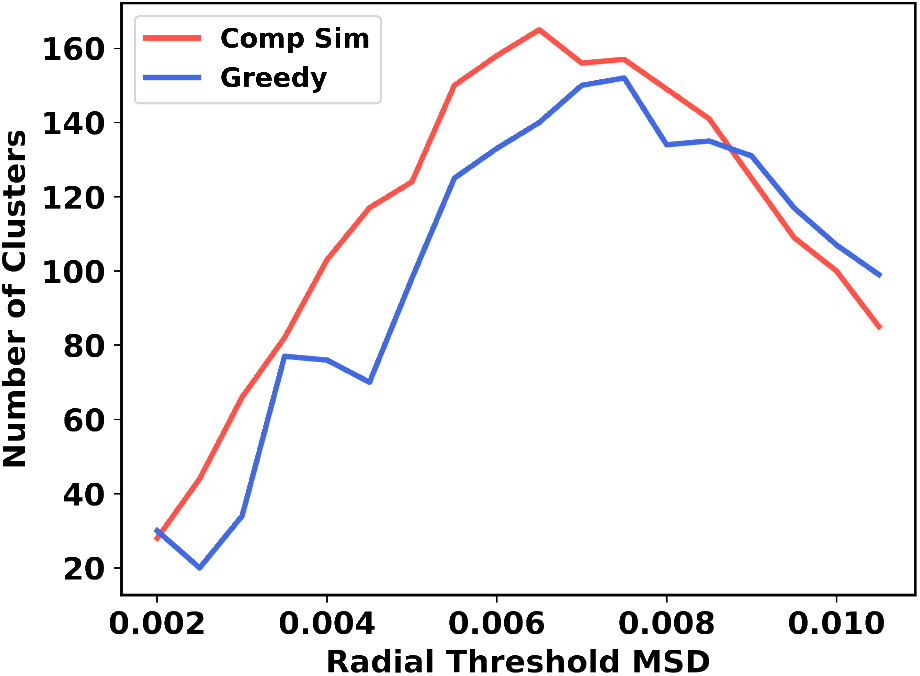
Screening of cluster numbers using the *k*-means++ vs comp sim seed selector.

##### Mean Squared Deviation

Although root-mean-square deviation (RMSD) is a standard similarity metric for its straightforward implementation and ease of interpretation, calculating the full pairwise RMSD matrix demands *O*(*N* ^2^) time and memory complexity. To alleviate these issues, we incorporate the notion of *n*-ary comparisons, that allow processing *N* objects with only *O(N)* cost. We have shown that this approach can be transformative in drug-design studies, as well as in the analysis of MD simulations, in which case we use the Mean Squared Deviation (MSD) as a way to quantify the separation between multiple conformation/frames. To calculate the MSD between a group of frames the column-wise sum of the normalized *N* × *D* matrix is calculated, in which we can rank the similarity of frames in the set.

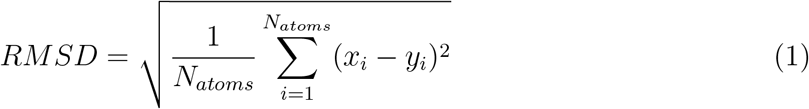

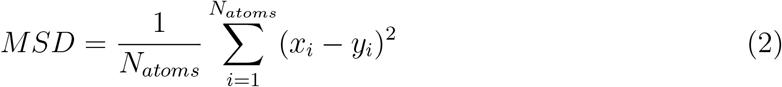

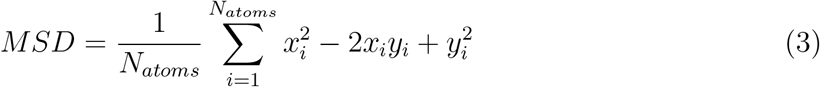

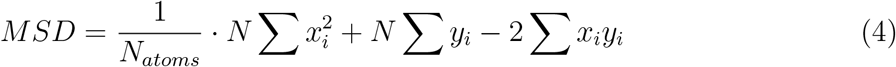

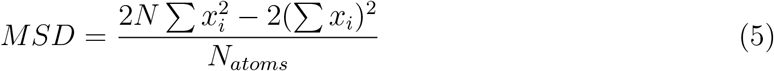

### Extended Quality Features

The defining advantage of eQual is that it does not require computing the pairwise RMSD matrix. Additionally, it provides an intuitive tie-breaking criterion that make the resulting method completely invariant to the order of the input frames, which makes it completely reproducible. Remember that although RTC can produce high-quality clusters, it is effectively an *O*(*N* ^2^) algorithm because every available configuration is proposing a cluster for every iteration. In addition, whenever there is a tie between densely populated clusters of the same size, the winner is varied because the winner is picked at random. We hope to improve on these shortcomings without compromising the final cluster accuracy.

#### Seed selection

Several seed selection methods are implemented in eQual, which are ultimately the key to turning this into a linear-scaling method. The two studied in this publication are complementary similarity using MSD and the *k*-means++ algorithm. Complementary similarity way stratifies the matrix of frames, allowing to easily distinguish between high- and low-density regions. After which, a diversity selection is performed to select the most representative, yet diverse frames as seeds for initializing eQUAL. The *k*-means++ algorithm uses a relatively similar intuition. ^23^ It finds the most different centroids in every iteration, which are spread according to a probability distribution function that tends to maximize their separation. (It is important to remark that any other *O(N)* clustering method could be used in this step, for example, *k*-means NANI, however, we decided to use *k*-means++ instead to make eQual as stand-alone as possible.) Both seed selection methods are similar in finding a diverse set of seeds for initializing the cluster formation. A critical advantage of complementary similarity is that it is deterministic because it aims for the lowest extended similarity between the current seeds and the candidate seed. ^30^ On the other hand, *k*-means++ results can vary due to its random nature of iterating through different centroids. However, this variability is more pronounced in the outlier sectors of the clusters, and since we are only interested in the cluster centroids, we did not observe appreciable variability in several eQual runs.

#### Tiebreaker criteria

After every seed proposes a cluster, the most populated cluster (within the given pre-selected threshold) is selected for that iteration. When there is more than one most populated cluster, a criterion is needed to break this tie. In Daura’s algorithm, he would take the first one in the list. However, this is not invariant to the order of the frames. In eQual, the MSD similarity for each proposed cluster is calculated and the proposed cluster with the lowest MSD will be selected for that iteration.

## Materials and Methods

### *N*-ary Continuous Similarity for Molecular Simulations

Only the coordinates of a trajectory are used and converted into a *N* × *D* matrix. *N* is the number of frames or the number of points that were sampled within a simulation time. *D* is three times the number of the selected atoms for (*x, y, z*) coordinates.^40^ It is then normalized between a [0, 1] interval to be compatible with the extended similarity indices which take in values in that range. The normalization is meant to agree with RMSD way of ranking. The minimum and maximum are taken over all coordinates.

### eQUAL Algorithm

Instead of an RMSD pairwise matrix, eQual only requires a *N* × *D* matrix and a radial threshold (given by the user as the only parameter in the procedure). The algorithm starts with a seed, determined either by the *k*-means++ algorithm or the medoids of the set. The seed is then removed from the matrix and the MSD of every frame to the seed is calculated. The frames that are within the threshold distance are added to the cluster. The clusters are then ranked by the number of frames in the cluster. The cluster with the most frames is the winner. In the case of a tie, the ultimate winner is the one with the highest extended similarity. The winner is then removed from the list and the process is repeated until there are no more frames left. A dictionary is returned with the cluster number as the key and the frames in the cluster as the value. An important point should be made about the use of *k*- means as a seed selector: typical applications of *k*-means need to identify an optimum value for *k*. However, this is not necessary for the eQual workflow, since we only need an estimate of the seed to grow the next cluster. For this, we just need to evaluate a few candidates. This is why in eQual, whenever we use *k*-means to find potential seeds, we just run *k*-means with a fixed value of *k* (typically, *k* = 5), and those 5 potential seeds are ranked as described above.

### Molecular Dynamics Simulation

The topology and trajectory files were taken from the github.com/LQCT/BitQT repo. The atom selection follows Daura et al, which is Lys2 to Asp11, with N, C*α*, C, O, and H atoms. The terminal and side chain residues were ignored to minimize noise in the clustering.

## Results and discussion

In the following we compare both variants of eQual: using *k*-means++ or complementary similarity to identify potential seed candidates. As expected, both methods produce largely equivalent results, with the *k*-means++ variant resulting in slightly more compact and well-separated clusters. Importantly, the eQual results are consistent with the RTC findings by Daura et al., indicating that the time and memory improvements in eQual come at no cost in the final quality of the clustering.

One of the key insights that we can gain from eQual is how the MSD threshold selected by the user ultimately affects the partitioning of the data. Every radial-based clustering (be it Taylor-Butina, BIRCH, BitBIRCH, RTC, etc.) is expected to follow the same general trend regarding the number of clusters vs. threshold tendency, with both the low- and large-threshold limits corresponding to low cluster counts (but with quite disparate overall partitions), with the general trend resembling an inverted parabola with usually a clear maximum in the number of clusters. At low distance threshold values the few clusters observed correspond to tightly-packed “hyper-spheres”, indicative of the centers of the high-density regions in the data. In this regime, it is to be expected that a substantial fraction of the conformations shows as singletons, so they are essentially considered “noise”. On the other hand, large distance thresholds result in virtually every point being assigned to a handful of very diffuse clusters. So, while no explicit outliers are found by the algorithm, the clusters formed in this region could be very heterogeneous, and thus of low quality. As shown in Fig. 3, eQual follows this same trends, with the *k*-means++ and complementary similarity variants having almost parallel behaviors across the range of MSD values explored.

**Figure 3.**
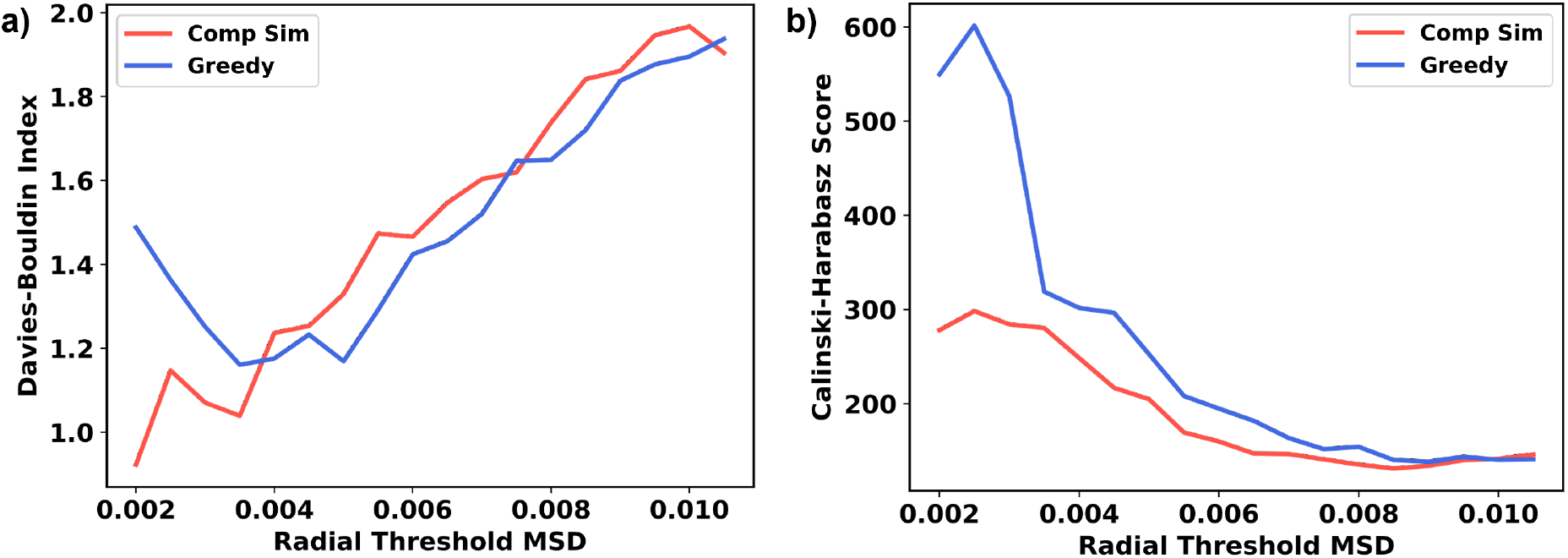
Screening of clustering scoring indices using the *k*-means++ vs `comp sim` seed selector.

The next obvious question to ask is then: which MSD thresholds one should use to analyze the data? In lieu of some pre-determined criterion (e.g., maybe we are just interested in studying changes up to a given maximum deformation from the cluster centers), this problem demands doing a scan of possible threshold values, and then using some clustering quality indicator to evaluate how well-separated is the data for each MSD. In Fig. 3, we show the results for the *k*-means++ and complementary similarity. Here, we decided to test two well-known clustering quality metrics: the Davies-Bouldin ^41^ and Calinski-Harabasz indices^42^ (DBI and CHI, respectively). These indices essentially quantify how compact and well-separated are the clusters, with the DBI slightly favoring the former, while the CHI tends to favor the latter. The traditional way of analyzing the results of these quality indices has been to look for the absolute minimum (DBI) or maximum (CHI) values, which tend to correspond to a better partitioning of the data. However, it has been noted before that these absolute comparisons might be biased to extreme situations, as can be seen in Fig. 3. Note how the DBI and CHI tend to favor the very low MSD regime, which is not surprising, since this leads to the smallest cluster sizes which are, by construction, more tightly-packed and with relatively sparser centroids. This is why we have proposed to complement the global analysis of the DBI and CHI values with a local analysis based on the relative variation of the indices relative to neighboring MSD thresholds. In short, the second derivative of the indices at each MSD value contains information on how stable is a given partition. In this context, a large (small) second derivative for the DBI (CHI) is preferred. These results are summarized in Table 1. Note how the DBI *k*-means++ analysis is quite consistent, with both the global and local analyses agreeing in that the 0.0035 and 0.0050 thresholds are favored. The DBI complementary similarity results also highlight 0.0035 as producing a stable segmentation of the frames, but the 0.0020 and 0.0060 thresholds indicate less robust results than the *k*-means++ seed selection. As observed in the case of *k*-means NANI, the CHI analysis is not as useful, since for this indicator the tendency to prefer very low cluster counts is the ultimate determining factor, as exemplified by the very low MSD thresholds in Table 1. This is why we suggest the use of the global and local DBI analysis as the preferred way to identify optimum clustering conditions. Finally, both for DBI and CHI Fig. 3 shows that the *k*-means++ seed selector tends to outperform the complementary similarity selection.

**Table 1:** Performance comparison of methods based on DBI and CHI metrics.

|  |  | Absolute value |  | 2nd deriv |  |
| --- | --- | --- | --- | --- | --- |
|  |  | Best | 2nd best | Best | 2nd best |
| DBI | Greedy | 0.0035 | 0.0050 | 0.0050 | 0.0035 |
|  | Comp_sim | 0.0020 | 0.0035 | 0.0035 | 0.0060 |
| CHI | Greedy | 0.0025 | 0.0020 | 0.0025 | 0.0080 |

This was expected, since the latter is a one-shot procedure (the potential seeds are selected once and are kept fixed), while the *k*-means++ procedure allows the initial candidate seeds to be optimized, by converging the *k*-means++ partitions at every step.

Having established the *k*-means++ as the more robust seed selector, we proceeded to study the concordance of the eQual/*k*-means++ results with those obtained through the classical RTC algorithm. First, Fig. 4 shows the population analyses for both the 0.0035 and 0.0050 MSD thresholds. Both thresholds lead to remarkably consistent populations for the top 10 denser clusters. In only one case (the 2nd most populated cluster) is the difference bigger than 1% (1.1%), while for the remaining clusters the differences are never bigger than 8 %, and in eight cases the differences are below 0.5%. As expected, the bigger MSD leads to clusters slightly bigger, but for both thresholds the top 10 clusters include ∼20% of the total conformations (19.8% for 0.0035 and 23% for 0.0050). These results compare very favorably with the RTC findings. Daura observed that there are 6 clusters with more than 100 conformations, ^39^ and we found 6 and 7 of those clusters for the 0.0035 and 0.0050 thresholds respectively. Moreover, all such clusters when MSD = 0.0035 account for 15.8% of the total population, and 19.6% in the case of MSD = 0.0050, which is very close to the 19% found by Daura.

**Figure 4.**
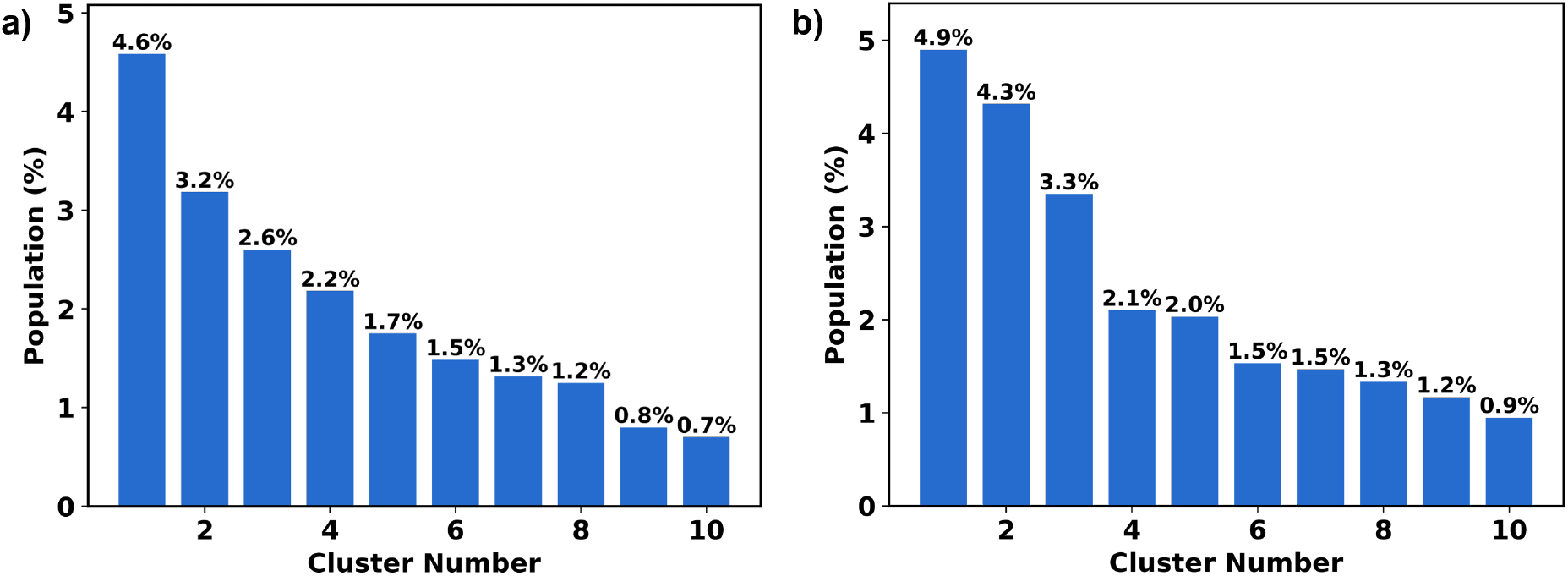
Using the *k*-means++ seed selector, **(a)** is the population of cluster found with the minimum DBI, **(b)** is the population of cluster found with maximum 2^nd^ derivative DBI.

The RMSD distributions (from all the conformations in a cluster to the cluster’s centroid) for the top 7 most populated clusters (those with ∼100 elements) also closely follows Daura’s results (Fig. 5). For MSD = 0.0035, all the curves are essentially centered around 0.5 Å, which aligns perfectly with Fig. 4 in Ref. 39. Even a greater MSD = 0.0050 shows very tight clusters, with the top 5 being closely packed around 0.5 Å, and only the lower-populated 6th and 7th clusters showing a displacement towards 1 Å and 1.5 Å, respectively. Yet again, this mirrors Daura’s observations in Figs. 4 and 5 in Ref. 39. Also notice the close match between the individual cluster populations reported by Daura in those figures and our results presented in Fig. 4.

**Figure 5.**
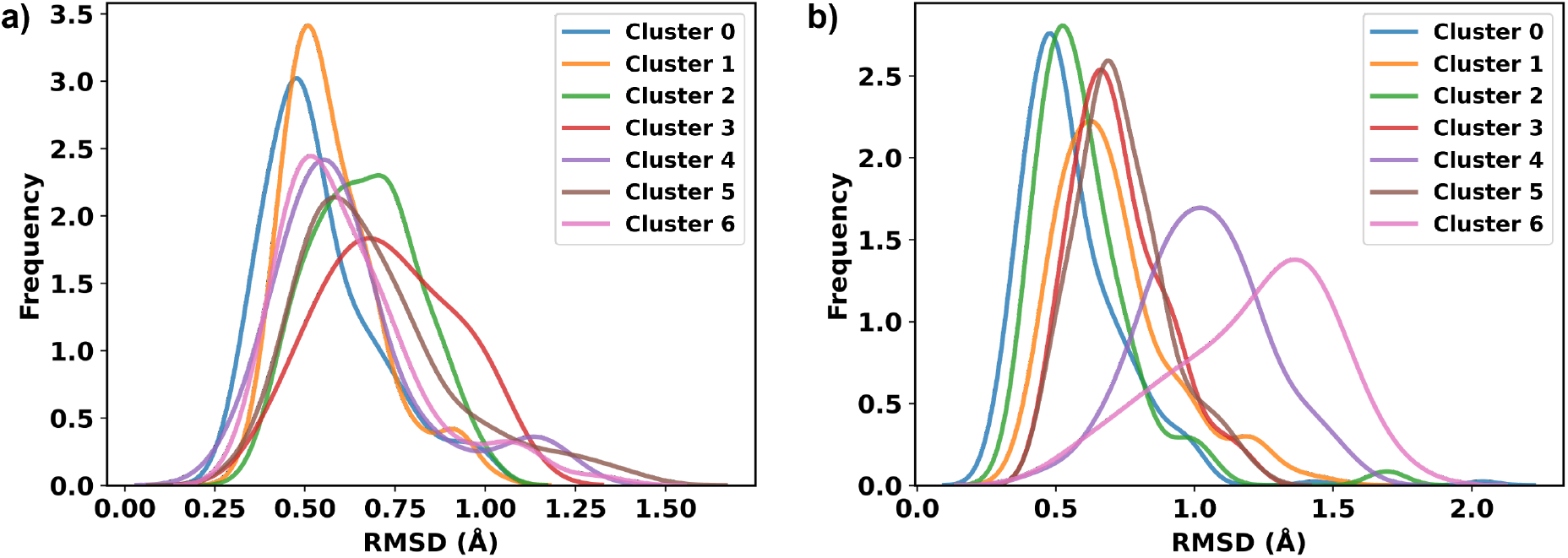
Structural overlaps *β*-heptapeptide shown for the eight top clusters. Clustering was conducted using the radial threshold with the maximum 2^nd^ derivative DBI (*MSD* = 0.005), *k*- means++ seed selector,.

Finally, Fig. 6 and 7 presents the main conformations found at thresholds 0.0035 and 0.0050. These conformations match those reported by Gonzalez-Aleman, ^26^ with the same characteristic “U”-, “S”-, and “scarf”-like patterns also observed in these cases. Once again, the overlaps of some conformations (shown in blue) with the cluster representative (medoid, shown in orange) highlight the ability of eQual of finding closely-packed clusters.

**Figure 6.**
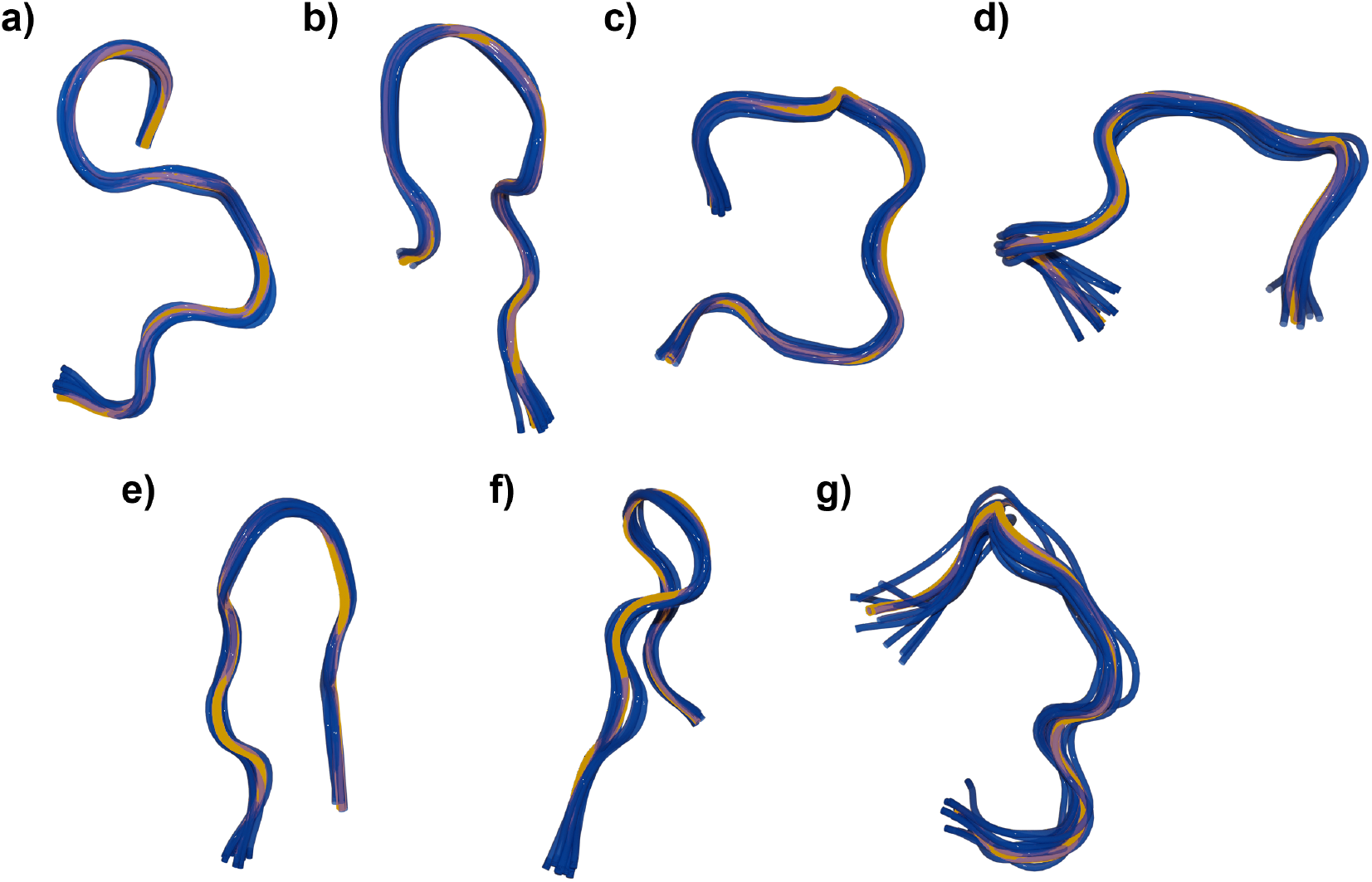
Structural overlaps *β*-heptapeptide shown for the seven top clusters. Clustering was performed using the radial threshold with the minimum DBI (*MSD* = 0.0035), *k*-means++ seed selector,.

**Figure 7.**
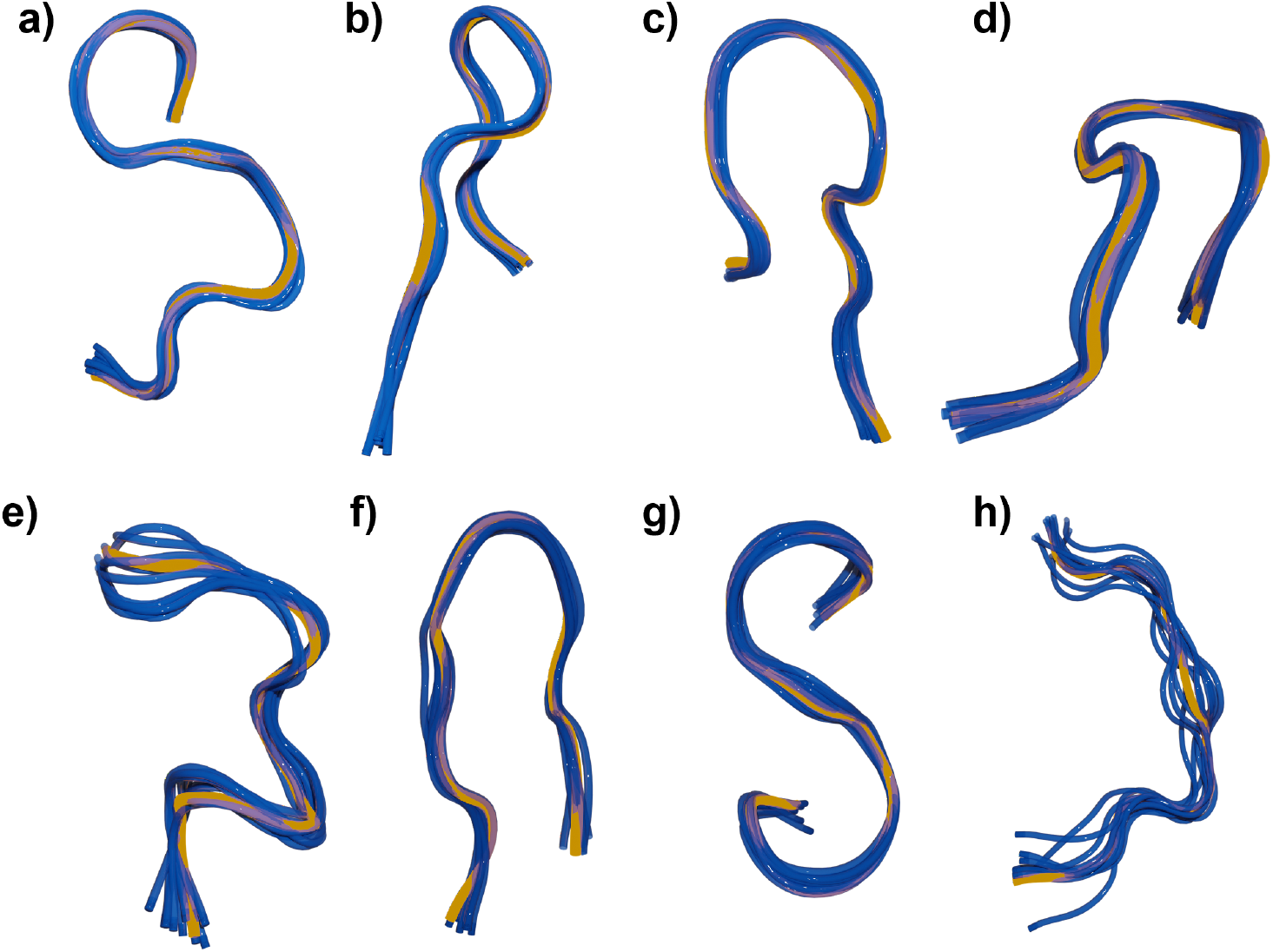
Structural overlaps *β*-heptapeptide shown for the eight top clusters. Clustering was conducted using the radial threshold with the maximum 2^nd^ derivative DBI (*MSD* = 0.005), *k*- means++ seed selector,.

## Conclusion

In summary, we have presented eQual, a radial threshold clustering algorithm that greatly improves on previous approaches by reducing the time and memory requirements, while not sacrificing the quality of the final clusters. eQual builds on the very intuitive notion of radial similarity to group related conformations, identifying promising starting seeds to grow the different clusters. The defining feature separating our approach from other algorithms in this family is that we do not have to explore every possible candidate starting point in order to select key conformations from dense regions. The two seed selection procedures proposed here (using complementary similarity, or just a few rounds of k-means++) produce largely equivalent results when it comes to number of clusters and their populations. As expected, the extra flexibility of the k-means++ initializer (that allows to relax the original selection of the frames) results in slightly better-separated and more compact clusters. Both approaches closely reproduce the RTC results, down to the populations of the denser clusters and the more representative conformations found by each method. In a sense, eQual bridges the gap between k-means and hierarchical approaches. On one hand, it has the same time and memory requirements of k-means, while avoiding what seems to be the biggest factor holding back the popularity of this method in the MD community: the need to pre-select an optimum value of k. While we have argued elsewhere that this is not really an issue, there is no denying that for most MD practitioners it is more natural to ask for clusters defined up to a given RMSD threshold from a given reference or representative structure. This is easily achievable in hierarchical methods, but these are plagued by highly inefficient time and (specially) memory management (in the absolute best case, a hierarchical method will be *O*(*N* ^2^), and even this needs to be coded very carefully). eQual, on the other hand, explicitly operates with a radial threshold, but without the need to pre-compute or store any pairwise distance matrices. eQual is part of the MDANCE package, and it is freely accessible at: https://github.com/mqcomplab/MDANCE.

## Acknowledgement

RAMQ and LC thank support from the National Institute of General Medical Sciences of the National Institutes of Health under award number R35GM150620.

## Notes

https://github.com/mqcomplab/MDANCE

### Competing Interest Statement

The authors have declared no competing interest.

## References

(1) Bowman, G. R.; Beauchamp, K. A.; Boxer, G.; Pande, V. S. Progress and challenges in the automated construction of Markov state models for full protein systems. The Journal of Chemical Physics 2009, 131, 124101.

(2) Friedrichs, M. S.; Eastman, P.; Vaidyanathan, V.; Houston, M.; Legrand, S.; Beberg, A. L.; Ensign, D. L. et al. Accelerating molecular dynamic simulation on graphics processing units. Journal of Computational Chemistry 2009, 30, 864–872.

(3) McGibbon, R. T.; Beauchamp, K. A.; Harrigan, M. P.; Klein, C.; Swails, J. M.; Hernández, C. X.; Schwantes, C. R. et al. MDTraj: a modern open library for the analysis of molecular dynamics trajectories. Biophysical journal 2015, 109, 1528–1532.

(4) Durrant, J. D.; McCammon, J. A. Molecular dynamics simulations and drug discovery. BMC Biology 2011, 9, 71.

(5) De Vivo, M.; Masetti, M.; Bottegoni, G.; Cavalli, A. Role of Molecular Dynamics and Related Methods in Drug Discovery. Journal of Medicinal Chemistry 2016, 59, 4035– 4061.

(6) Trozzi, F.; Wang, X.; Tao, P. UMAP as a Dimensionality Reduction Tool for Molecular Dynamics Simulations of Biomacromolecules: A Comparison Study. The Journal of Physical Chemistry B 2021, 125, 5022–5034.

(7) Melvin, R. L.; Godwin, R. C.; Xiao, J.; Thompson, W. G.; Berenhaut, K. S.; Salsbury, F. R. Uncovering Large-Scale Conformational Change in Molecular Dynamics without Prior Knowledge. Journal of Chemical Theory and Computation 2016, 12, 6130–6146.

(8) Tribello, G. A.; Gasparotto, P. Using Dimensionality Reduction to Analyze Protein Trajectories. Frontiers in Molecular Biosciences 2019, 6, 46.

(9) Doerr, S.; Ariz-Extreme, I.; Harvey, M. J.; Fabritiis, G. D. Dimensionality reduction methods for molecular simulations. 2017; http://arxiv.org/abs/1710.10629, x1710.10629 [stat].

(10) Shao, J.; Tanner, S. W.; Thompson, N.; Cheatham, T. E. Clustering Molecular Dynamics Trajectories: 1. Characterizing the Performance of Different Clustering Algorithms. Journal of Chemical Theory and Computation 2007, 3, 2312–2334.

(11) Shortle, D.; Simons, K. T.; Baker, D. Clustering of low-energy conformations near the native structures of small proteins. Proceedings of the National Academy of Sciences 1998, 95, 11158–11162.

(12) Daura, X.; Gunsteren, W. F. v.; Mark, A. E. Folding–unfolding thermodynamics of a beta-heptapeptide from equilibrium simulations. Proteins: Structure, Function, and Bioinformatics 1999, 34, 269–280.

(13) Daura, X.; Gademann, K.; Jaun, B.; Seebach, D.; Van Gunsteren, W. F.; Mark, A. E. Peptide Folding: When Simulation Meets Experiment. Angewandte Chemie International Edition 1999, 38, 236–240.

(14) Lemke, O.; Keller, B. Common Nearest Neighbor Clustering—A Benchmark. Algorithms 2018, 11, 19.

(15) Rodriguez, A.; Laio, A. Clustering by fast search and find of density peaks. Science 2014, 344, 1492–1496.

(16) Wang, S.; Chang, T.-H.; Cui, Y.; Pang, J.-S. Clustering by orthogonal NMF model and non-convex penalty optimization. IEEE Transactions on Signal Processing 2021, 69, 5273–5288.

(17) De Paris, R.; Quevedo, C. V.; Ruiz, D. D. A.; Norberto de Souza, O. An Effective Approach for Clustering InhA Molecular Dynamics Trajectory Using Substrate-Binding Cavity Features. PLOS ONE 2015, 10, e0133172.

(18) Keller, B.; Daura, X.; van Gunsteren, W. F. Comparing geometric and kinetic cluster algorithms for molecular simulation data. The Journal of Chemical Physics 2010, 132, 074110.

(19) Glielmo, A.; Husic, B. E.; Rodriguez, A.; Clementi, C.; Noé, F.; Laio, A. Unsupervised Learning Methods for Molecular Simulation Data. Chemical Reviews 2021, 121, 9722– 9758.

(20) Konovalov, K. A.; Unarta, I. C.; Cao, S.; Goonetilleke, E. C.; Huang, X. Markov State Models to Study the Functional Dynamics of Proteins in the Wake of Machine Learning. JACS Au 2021, 1, 1330–1341.

(21) Husic, B. E.; Pande, V. S. Markov State Models: From an Art to a Science. Journal of the American Chemical Society 2018, 140, 2386–2396.

(22) Yang, Y. I.; Shao, Q.; Zhang, J.; Yang, L.; Gao, Y. Q. Enhanced sampling in molecular dynamics. The Journal of Chemical Physics 2019, 151, 070902.

(23) Arthur, D.; Vassilvitskii, S. k-means++: the advantages of careful seeding. Proceedings of the eighteenth annual ACM-SIAM symposium on discrete algorithms. USA, 2007; pp 1027–1035.

(24) Kaufman, L.; Rousseeuw, P. J. Finding groups in data: An introduction to cluster analysis.; John Wiley, 1990.

(25) Butina, D. Unsupervised Data Base Clustering Based on Daylight’s Fingerprint and Tanimoto Similarity: A Fast and Automated Way To Cluster Small and Large Data Sets. Journal of Chemical Information and Computer Sciences 1999, 39, 747–750.

(26) González-Alemán, R.; Hernández-Castillo, D.; Caballero, J.; Montero-Cabrera, L. A. Quality Threshold Clustering of Molecular Dynamics: A Word of Caution. Journal of Chemical Information and Modeling 2020, 60, 467–472.

(27) Danalis, A.; McCurdy, C.; Vetter, J. S. Efficient Quality Threshold Clustering for Parallel Architectures. 2012 IEEE 26th International Parallel and Distributed Processing Symposium. Shanghai, China, 2012; pp 1068–1079.

(28) Chen, L.; Roe, D. R.; Kochert, M.; Simmerling, C.; Miranda-Quintana, R. A. k-Means NANI: An Improved Clustering Algorithm for Molecular Dynamics Simulations. Journal of Chemical Theory and Computation 2024, 20, 5583–5597.

(29) López-Pérez, K.; Kim, T. D.; Miranda-Quintana, R. A. iSIM: instant similarity. Digital Discovery 2024, 3, 1160–1171.

(30) Miranda-Quintana, R. A.; Rácz, A.; Bajusz, D.; Héberger, K. Extended similarity indices: the benefits of comparing more than two objects simultaneously. Part 2: speed, consistency, diversity selection. Journal of Cheminformatics 2021, 13, 33.

(31) Miranda-Quintana, R. A.; Bajusz, D.; Rácz, A.; Héberger, K. Extended similarity indices: the benefits of comparing more than two objects simultaneously. Part 1: Theory and characteristics†. Journal of Cheminformatics 2021, 13, 32.

(32) Rácz, A.; Dunn, T. B.; Bajusz, D.; Kim, T. D.; Miranda-Quintana, R. A.; Héberger, K. Extended continuous similarity indices: theory and application for QSAR descriptor selection. Journal of Computer-Aided Molecular Design 2022, 36, 157–173.

(33) Chang, L.; Perez, A.; Miranda-Quintana, R. A. Improving the analysis of biological ensembles through extended similarity measures. Physical Chemistry Chemical Physics 2022, 24, 444–451.

(34) López-Pérez, K.; Jung, V.; Chen, L.; Huddleston, K.; Miranda-Quintana, R. A. Efficient clustering of large molecular libraries. 2024; http://biorxiv.org/lookup/doi/10.1101/2024.08.10.607459.

(35) Dunn, T. B.; Seabra, G. M.; Kim, T. D.; Juárez-Mercado, K. E.; Li, C.; Medina-Franco, J. L.; Miranda-Quintana, R. A. Diversity and Chemical Library Networks of Large Data Sets. Journal of Chemical Information and Modeling 2022, 62, 2186–2201.

(36) López-Pérez, K.; Avellaneda-Tamayo, J. F.; Chen, L.; López-López, E.; Juárez-Mercado, K. E.; Medina-Franco, J. L.; Miranda-Quintana, R. A. Molecular similarity: Theory, applications, and perspectives. Artificial Intelligence Chemistry 2024, 2, 100077.

(37) Ellin, N. R.; Guo, Y.; Miranda-Quintana, R. A.; Prentice, B. M. Extended similarity methods for efficient data mining in imaging mass spectrometry. Digital Discovery 2024, 3, 805–817.

(38) Chen, L.; Mondal, A.; Perez, A.; Miranda-Quintana, R. A. Protein Retrieval via Integrative Molecular Ensembles (PRIME) through Extended Similarity Indices. Journal of Chemical Theory and Computation 2024, Publisher: American Chemical Society.

(39) Daura, X.; Conchillo-Solé, O. On Quality Thresholds for the Clustering of Molecular Structures. Journal of Chemical Information and Modeling 2022, 62, 5738–5745.

(40) Rácz, A.; Mihalovits, L. M.; Bajusz, D.; Héberger, K.; Miranda-Quintana, R. A. Molecular Dynamics Simulations and Diversity Selection by Extended Continuous Similarity Indices. Journal of Chemical Information and Modeling 2022, 62, 3415–3425.

(41) Davies, D. L.; Bouldin, D. W. A Cluster Separation Measure. IEEE Transactions on Pattern Analysis and Machine Intelligence 1979, PAMI-1, 224–227.

(42) Caliński, T.; Harabasz, J. A dendrite method for cluster analysis. Communications in Statistics 1974, 3, 1–27.

